# Integrating Single-Molecule Sequencing and Deep Learning to Predict Haplotype-Specific 3D Chromatin Organization in a Mendelian Condition

**DOI:** 10.1101/2025.02.26.640261

**Authors:** Danilo Dubocanin, Anna Kalygina, J. Matthew Franklin, Cy Chittenden, Mitchell R Vollger, Shane Neph, Andrew B Stergachis, Nicolas Altemose

**Affiliations:** Department of Genetics, School of Medicine, Stanford University, Palo Alto, CA, USA; Department of Biology, University of Oxford, Oxford, UK; Division of Medical Genetics, Dept. of Medicine, University of Washington, Seattle, WA, USA; Department of Genome Sciences, University of Washington, Seattle, WA, USA; Brotman Baty Institute for Precision Medicine, Seattle, WA, USA; Chan Zuckerberg Biohub – San Francisco, San Francisco, CA, USA

## Abstract

The three-dimensional (3D) architecture of the genome plays a crucial role in gene regulation and various human diseases. Short-read sequencing methods for measuring 3D genome organization are powerful, but they lack the ability to resolve individual human haplotypes or structurally complex regions. To address this, we present FiberFold, a deep learning model that combines convolutional neural networks and transformer architectures to accurately predict cell-type-specific and haplotype-specific 3D genome organization using multi-omic data from a single, long-read sequencing assay, Fiber-seq. By applying FiberFold to a cell line with allelic X-inactivation, we show that Topologically Associated Domains (TADs) are attenuated on the inactive chrX. Furthermore, FiberFold predicts significant changes to TADs surrounding a 13;X balanced translocation in a patient with a rare Mendelian disease. FiberFold showcases the power of integrating long-read epigenomic sequencing with deep learning tools to investigate fundamental chromatin biology as well as the molecular basis of human disease.

## INTRODUCTION

Given the need to compact two meters of DNA into each cell’s nucleus, humans, like all complex organisms, have evolved a set of mechanisms to organize their genome in 3D space ^1^. There is ever-increasing evidence linking 3D genome organization to fundamental cellular processes and various diseases ^2–9^. To explore 3D genome structure, various short-read sequencing-based methods like Hi-C and Micro-C have been developed ^10,11^. Similarly, techniques like ChIP-seq, ATAC-seq, and bisulfite sequencing have been developed to probe distinct epigenetic features ^12–15^. Building a complete understanding of the genomic regulatory machinery in different contexts necessitates the use of multiple short-read sequencing techniques, exacerbating experimental costs. Furthermore, these short-read approaches are fundamentally limited in their inability to resolve individual haplotypes, repetitive genomic regions, structural variants, and the combinatorial interactions between regulatory machinery.

Recently, multiple long-read, single-molecule sequencing methods including Fiber-seq ^16^, SMAC-seq ^17^, and DiMeLo-seq ^18^ have emerged, addressing some of the limitations of traditional short-read assays for probing chromatin accessibility and protein-DNA interactions ^19–24^ . For example, Fiber-seq has been used to examine the chromatin state in previously unmappable regions of the genome, such as centromeres and telomeres^25,26^, to generate haplotype-specific maps of genome regulation^27^, to study phenomena like transcription at the single-molecule level^28,29^, and to dissect disease mechanisms ^30,31^. Other methods that utilize long-read sequencing to capture 3D genome organization, such as PoreC ^32^, also provide rich multi-omic information, but they still require large amounts of input material and rely on the concatenation of shorter DNA fragments (< 500 bp), which limits their ability to measure 3D genome organization in structurally variable regions or on specific haplotypes. Furthermore, like other Hi-C-like methods, these approaches require super-linearly deeper sequencing in order to improve the resolution and sensitivity of their 3D genome structure maps^33,34^.

In parallel, recent advances in machine learning have enabled the development of frameworks to predict 3D genome organization directly from genome sequence ^35–37^. While these models can accurately reconstruct 3D genome architecture in the cell types used for model training, they are limited in their cross cell-type generalizability. The C.Origami framework was developed to integrate chromatin accessibility and CTCF binding information alongside genome sequence data to reconstruct de novo Hi-C maps, closely matching experimental Hi-C results ^38^. While powerful, this approach still requires multiple experiments and cannot provide haplotype-phased 3D genome structure information.

Additionally, concerns have been raised that sequence-to-function models might learn biologically implausible patterns from training data, further limiting the generalizability of sequence-based models ^39–42^. Recently, there has been progress in developing sequence-free models that may outperform traditional, sequence-based models ^43,44^.

Fiber-seq, with its ability to capture chromatin accessibility, CpG methylation, and CTCF binding in a single assay, presents an opportunity to streamline the generation of 3D genomic structure information. We hypothesized that by developing deep learning models with Fiber-seq data alone, we could accurately predict cell-type-specific Hi-C maps. Here we introduce FiberFold, a method that integrates chromatin accessibility, CTCF binding, and CpG methylation data from Fiber-seq alone to predict cell-type-specific and haplotype-specific 3D genome interactions from a single assay. We demonstrate that FiberFold accurately recapitulates 3D genome structure as captured by Hi-C within and across cell-types. We further utilize FiberFold to demonstrate that complex 3D topologies are attenuated on the inactive X chromosome. Finally, we showcase the clinical utility of FiberFold by predicting changes in 3D genome organization caused by an autosome-X chromosomal translocation in a patient with a Mendelian condition.

We anticipate that FiberFold will help to accelerate studies of genome regulation by enabling the rapid analysis of 3D genome organization in conjunction with chromatin accessibility, CTCF binding, CpG methylation, and underlying genetic variation. We believe that this is the first demonstration of the quantification of these 5 features of our individual genomes in one assay, and we hope that it will further increase the utility of single-molecule sequencing methods like Fiber-seq.

## RESULTS

### FiberFold Architecture, Training, and Evaluation

The FiberFold model architecture builds upon the foundational framework of C.Origami, with several important modifications to enhance performance and generalizability. Notably, FiberFold was designed to exclude the underlying DNA sequence from its training inputs, a deliberate choice aimed at improving the model’s ability to generalize across different haplotypes and cell types. Instead of relying on sequence information, FiberFold integrates three distinct feature tracks derived entirely from Fiber-seq data: chromatin actuation, which is measured using Fiber-seq Inferred Regulatory Elements (FIREs), CpG methylation (mCpG), and CTCF footprinting scores (CTCF FP). FIREs have been extensively profiled and have been shown to recapitulate signals captured by DNaseI-seq and ATAC-seq, and we show that CTCF footprinting with Fiber-seq is consistent with CTCF ChIP-seq signal (Fig. 1A,B, Supp. Fig. 1A,B) ^16,27,31^. We also included the directionality of CTCF motifs, given their functional relevance in determining 3D genomic structure ^45–48^. FiberFold takes in these features in a 2,097,152 bp window and applies consecutive 1D residual-convolutions to compress the information in a latent space (8,192 bp resolution) that is then fed into a transformer, which learns relational long-range interactions. The output of the transformer is then fed into consecutive 2-D dilated residual convolutions to output a predicted Hi-C map at 10 kb resolution (Fig. 1C,D).

**Figure 1:**
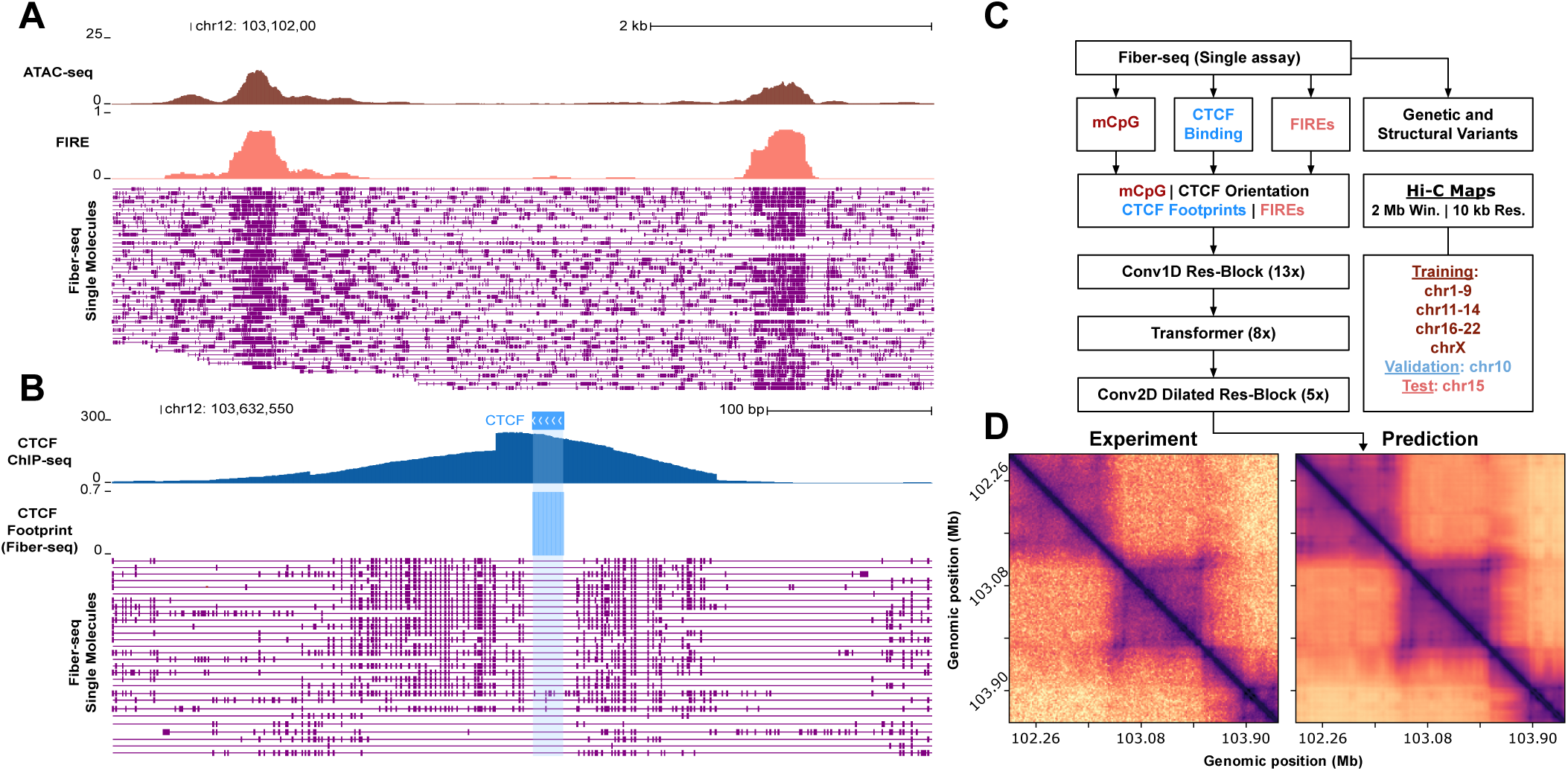
FiberFold Overview. **A,** Genome browser track depicting depicting ATAC-seq, Fiber-seq Inferred Regulatory elements (FiREs), and accompanying Fiber-seq single molecules, represented by purple lines, m6A on each molecule represented as box along line. **B**, Genome browser track depicting CTCF ChIP-seq, CTCF footprint score from Fiber-seq, and accompanying Fiber-seq single molecules. **C,** Framework and model architecture for FiberFold. **D,** Example prediction for genomic locus encompassing regions in **A,B.**

We performed training using Fiber-seq and Hi-C data from the extensively profiled human lymphoblast cell line GM12878 ^31^. FiberFold was trained to predict Hi-C maps by minimizing the mean squared error between experimentally derived and model-predicted maps at 10 kb resolution. We split the genome into 2 Mb windows, sliding across the genome with a 36 kb window with a randomized shift parameter applied to these windows to increase training data diversity. We did not include regions overlapping ENCODE Blacklist region s or regions with significant coverage gaps (Methods). For model evaluation, chromosomes were split into training, validation (chr10), and test (chr15) sets.

FiberFold’s performance on the test chromosome exhibited strong concordance with experimental Hi-C maps in map-to-map comparisons of the predicted bin values. (median Pearson r = 0.938, median Spearman r = 0.906, Fig. 2A,B). We also observed strong pixel-wise concordance on test-chromosome predictions across distances (Pearson r = 0.932, Fig. 2B,C, Supp. Fig. 2 A).

**Figure 2:**
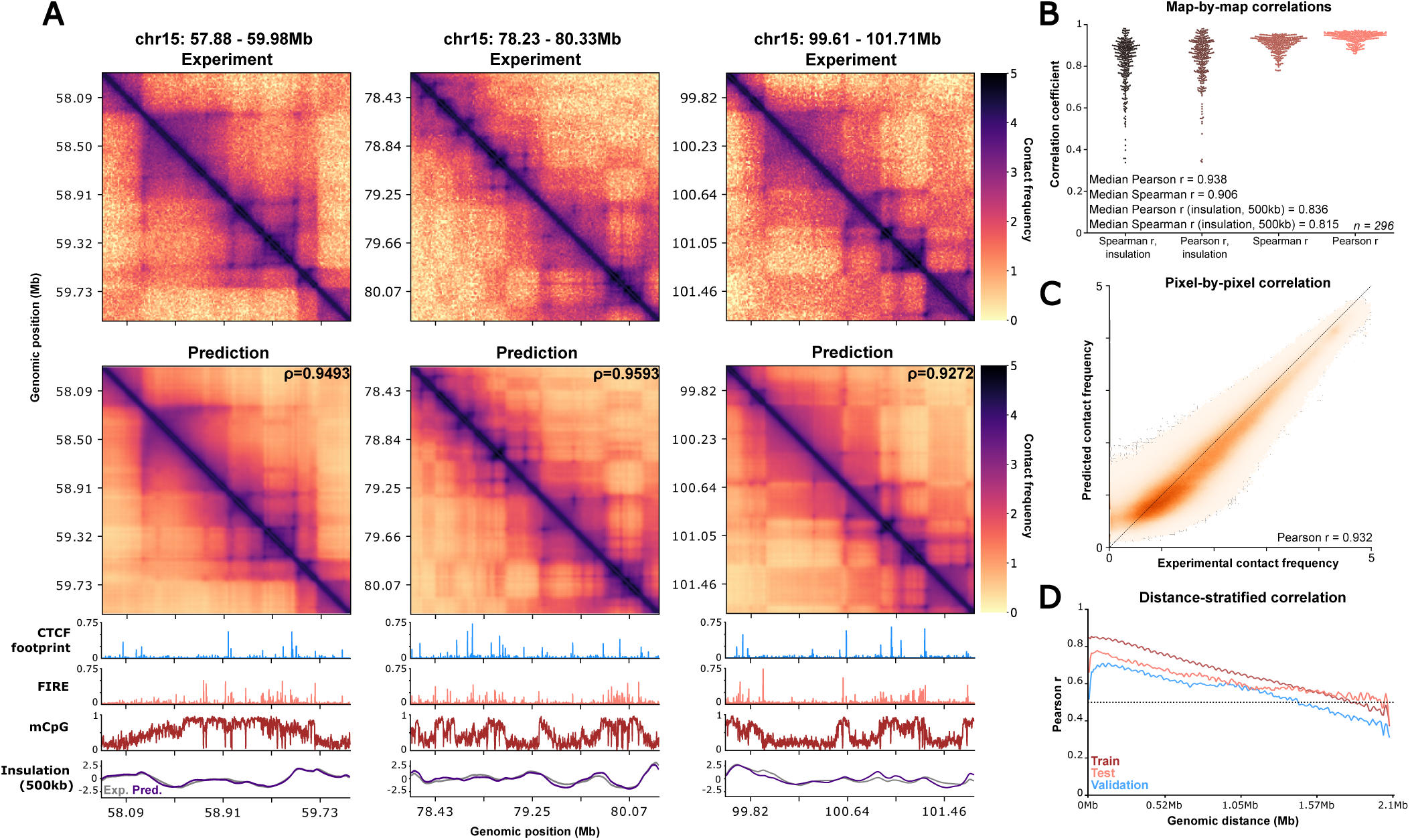
FiberFold predicts Hi-C maps accurately in GM12878. **A,** Experimental (top) and Predicted (middle) Hi-C maps for held-out test chromosome 15 in GM12878. Maps are natural-log scaled. Upper right corner of predicted map displays Pearson R between real and predicted maps. Below Predictions are input features used for the model and associated insulation for each map (radius = 500 kb). **B,** Distribution of various correlations for comparisons between real and predicted Hi-C maps for held-out chromosome 15. **C,** Pixel-by-pixel correlation for real and predicted values across chr15, not including contacts <16.4kb away. **D**, Distance stratified pixel-by-pixel correlation across the train, test, and validation sets.

We observed a significant positive correlation between CTCF footprinting scores and predicted insulation strength (*r*^2^ = .755, *p* = 0.0011, Supp. Fig. 2B), consistent with prior evidence that CTCF-mediated loop-extrusion is the major primary contributor to chromatin compartmentalization at this scale ^49–56^. Furthermore, there was a monotonic relationship between CTCF footprinting scores and predicted insulation score only at sites with high Rad21 ChIP-seq occupancy (Supp. Fig. 2C) ^57–59^, consistent with cohesin-dependent mechanistic principles of loop-extrusion^1,60,61^.

### FiberFold generalizes to a separate human cell type

3D genome structure tends to be highly conserved across diverse cell types ^55^. However, cell types can exhibit unique 3D genomic interactions that are biologically relevant, such as in cancers ^5,6,62,63^ and in both common and rare diseases ^7,64,65^. To assess the predictive capacity of FiberFold in a different cell type, we applied our GM12878-trained model to predict Hi-C maps using Fiber-seq data from a chronic myelogenous leukemia cell line (K562). Although K562 and GM12878 share a common lymphoid lineage differentiation pathway, previous work modeling cell-type specific contacts demonstrates that the 3D genomic organization between these two cell lines is as divergent as that observed between GM12878 and stem cells (H1 hESCs) ^38^. Our model maintained high predictive accuracy in K562 in both map-to-map comparisons (median ⍴ = 0.930, median ⍴s = 0.902) as well as pixel-to-pixel comparisons across stratified distances (⍴ = 0.871, Fig. 3A-C, Supp. Fig. 3 A-C) .

Furthermore, FiberFold successfully identified differential 3D chromatin organization patterns between GM12878 and K562, highlighting its capacity to predict cell-type-specific 3D chromatin architectures using Fiber-seq data alone (Fig. 3A, Supp. Fig. 3C).

**Figure 3:**
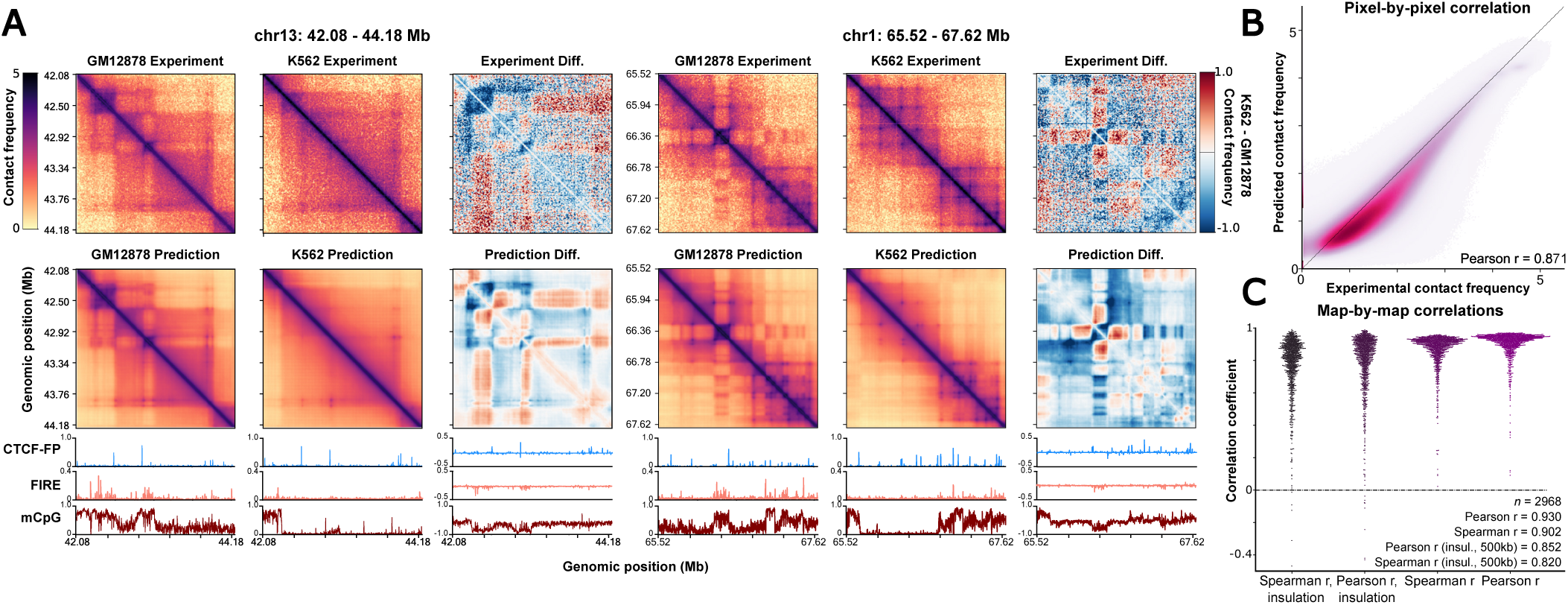
FiberFold predictions in a *de novo* cell line, K562. **A,** Experimental and Predicted Hi-C maps for 2 genomic loci for a held out cell type, K562 (middle), training cell type, GM12878 (left), and the difference between the maps (K562 - GM12878, right). Maps are natural-log scaled. Input features used for model predictions (left, middle), and the difference between features (right) are displayed below the contact maps. **B**, Pixel-by-pixel correlation across all chromosomes in K562. **C,** Distribution of various correlations for comparisons between real and predicted Hi-C maps for held-out cell type K562.

### Resolution of haplotype-selective autosomal 3D genomic structure with FiberFold

Human somatic cells contain two complete copies of the genome (haplotypes), and resolving the differences between how these two copies fold within a cell is necessary for understanding the effects of genetic variation. Current short-read technologies are poorly suited for comprehensively measuring haplotype-specific 3D genomic features in humans owing to the paucity of heterozygous sites across the genome (*e.g.,* less than 10% of Hi-C reads are typically assigned to a specific haplotype ^55^). As long-read sequencing data can be accurately haplotype phased, we reasoned that using haplotype-resolved Fiber-seq data with FiberFold would enable predictions of haplotype-selective 3D chromatin interactions. To test this, we haplotype-phased our GM12878 Fiber-seq data using parental short-read sequencing data, and applied FiberFold separately to each of the two haplotypes. We then evaluated our haplotype-specific FiberFold data along the GM12878 X chromosome, as these GM12878 cells are known to contain allelic skewing of X chromosome inactivation (XCI) (*i.e.,* the active X [Xa] is the paternal haplotype, and the inactive X [Xi] is the maternal haplotype) ^27^ (Fig. 4A).

Genome-wide, we observed that the averaged FiberFold-predicted Hi-C maps across both haplotypes correlated well with the experimentally derived Hi-C data (autosome Pearson r = 0.938, chrX Pearson r = 0.91) consistent with Hi-C experimental data measuring the averaged 3D genomic structure across both haplotypes (Supp. Fig. 4A). However, these haplotype-specific Hi-C maps enabled us to identify distinct genomic locations that diverged in their predicted 3D structure between the paternal and maternal haplotypes (Fig. 4B, Supp. Fig. 4C). Notably, chromosome X showed the highest haplotype-specific differences in predicted Hi-C maps (ind. t-test *p* < 1x10^-16^, Fig. 4B-D, Supp. Fig. 4B-D), indicating that FiberFold may be uncovering differences in the 3D structure of the active and inactive X chromosome. Specifically, the predominant alteration we observed in the predicted Hi-C maps along the Xi was widespread attenuation in sub-megabase scale TADs and distal contact formation (Fig. 4C). Stratification of haplotype-specific predicted Hi-C pixel values based on their matched experimental true value demonstrated that in chrX, the Xa (*i.e.,* paternal haplotype) disproportionately contributes to the upper and lower extrema, while this relationship is absent in autosomes (Supp. Fig. 4B).

Overall, our findings indicate that the human Xi and Xa likely adopt distinct 3D chromatin interactions, which is characterized by TAD attenuation along the Xi, a pattern that has previously only been observable using highly genetically divergent mouse crosses ^64–69^. Importantly, this demonstrates the potential of FiberFold for enabling the comprehensive study of chrX folding in humans as opposed to mice.

**Figure 4:**
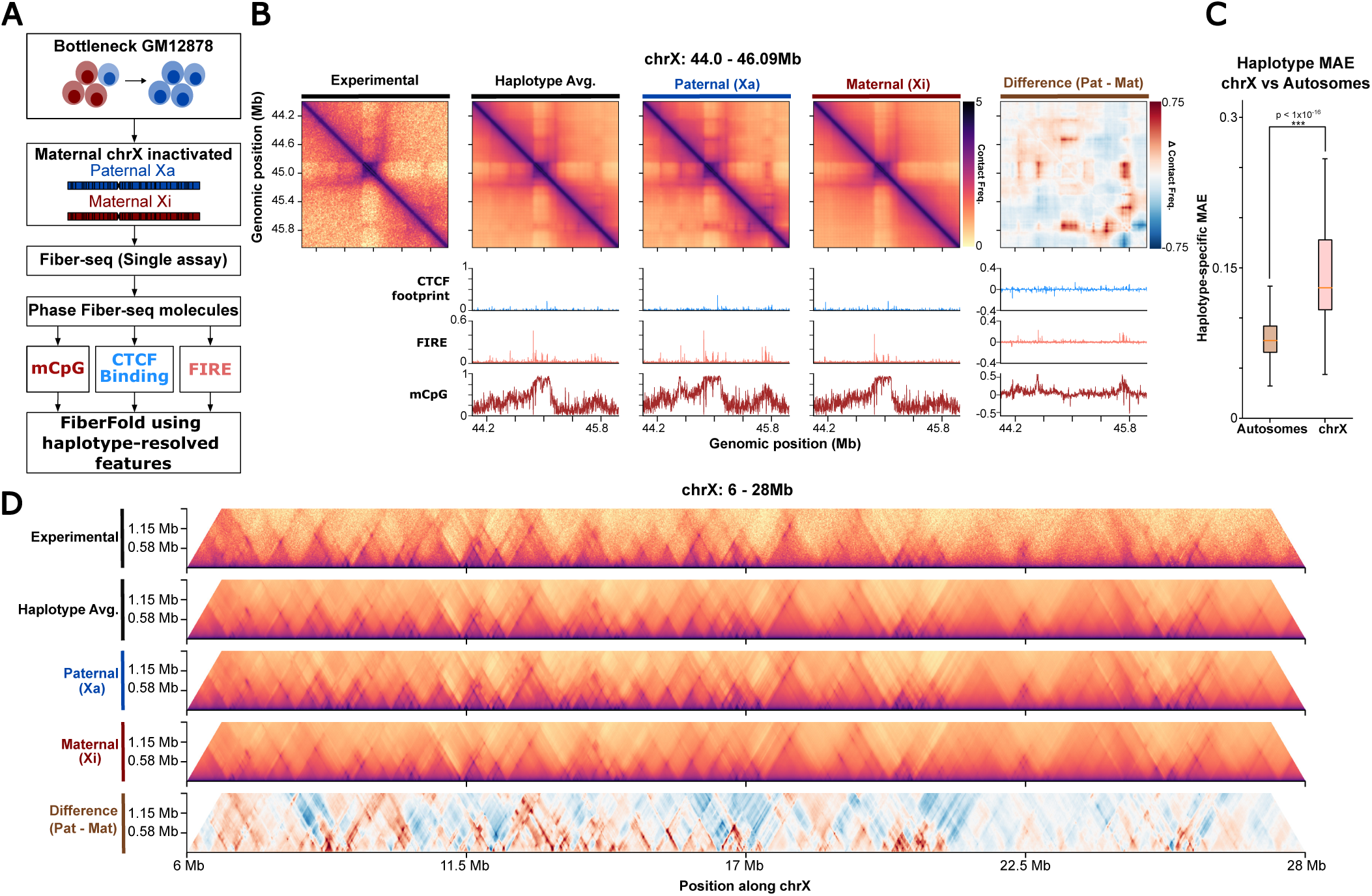
FiberFold enables haplotype specific predictions in a bottlenecked GM12878 line. **A**, Overview of process for predicting haplotype-specific Hi-C maps. **B** Locus from chrX displaying haplotype-specific Hi-C differences on chrX. From left: first map represents experimental Hi-C data; second map represents averaged Hi-C map across the haplotypes, and unphased input features; third map represents paternal predicted Hi-C map and paternal input features; fourth map represents maternal predicted Hi-C map and maternal input features; fifth map represents the difference in Hi-C maps and input features (paternal - maternal). **C,** Mean Absolute Error between haplotype-specific predictions, on autosomes or chrX (ind. t-test). **D,** (From top) Experimental, averaged across haplotypes, paternal, maternal, and differential Hi-C maps across a 22 Mb segment of chrX.

### Predicting 3D genome structure in a X;13 balanced translocation causing a Mendelian condition

To test the ability of FiberFold to resolve haplotype-specific 3D chromatin interactions that are clinically impactful, we next applied FiberFold to Fiber-seq data from a patient-derived cell line that harbors a X;13 balanced translocation (46,XX,t(X;13)(p22.1;q14.1)) (UDN318336) that was previously shown to cause a novel Mendelian condition associated with the dysregulation of multiple genes ^31^. The translocation resulted in a paternal haplotype with a 50 Mb derivative chromosome 13 (der(13)), which includes 13p, part of 13q, and an inverted Xp, as well as a 210 Mb derivative chromosome X (der(X)), encompassing Xq, part of Xp, and an inverted 13q ^31^ (Figure 5A). Furthermore, as is typical in X-autosome balanced translocations, the intact chrX is predominately subjected to XCI, resulting in der(X) remaining active ^31,66,67^.

Using Fiber-seq data from patient-derived retinal organoids with this translocation, and parental short-read sequencing data, we generated a haplotype-phased, contig-level assembly with N50 values of 82.3 Mb and 92.5 Mb for the paternal and maternal haplotypes, respectively. Importantly, we assembled 35 Mb+ contigs that spanned the translocation breakpoints (der(13) = 35.6 Mb, der(X) = 45.4 Mb, chrX = 93.1 Mb, chr13 = 87.2 Mb). Using this donor-specific assembly (DSA), we were able to accurately map the Fiber-seq reads back onto these contigs and generate the regulatory feature inputs for FiberFold in a haplotype-aware manner (Fig. 5A).

We observed a striking difference in the organization of the predicted 3D chromatin architecture in der(X) (*i.e.,* Xa) compared to the homologous portion of the intact chrX (*i.e.,* Xi). The predicted Hi-C maps on the intact chrX (*i.e.,* Xi) suggest the formation of large, moderately compacted interaction compartments, while predictions on der(X) indicate the formation of numerous sub-megabase TADs (Fig. 5A). The difference in 3D chromatin architecture between the homologous sequences of der(X) and intact chrX forms a distinctive ‘sawtooth’ pattern, indicating a significant and preferential formation of smaller TADs on the active der(X) relative to the inactive intact chrX, or a pronounced attenuation of TADs on the inactive intact chrX, consistent with our findings along Xi in GM12878 cells (Fig. 5A-B, Fig. 4D).

**Figure 5:**
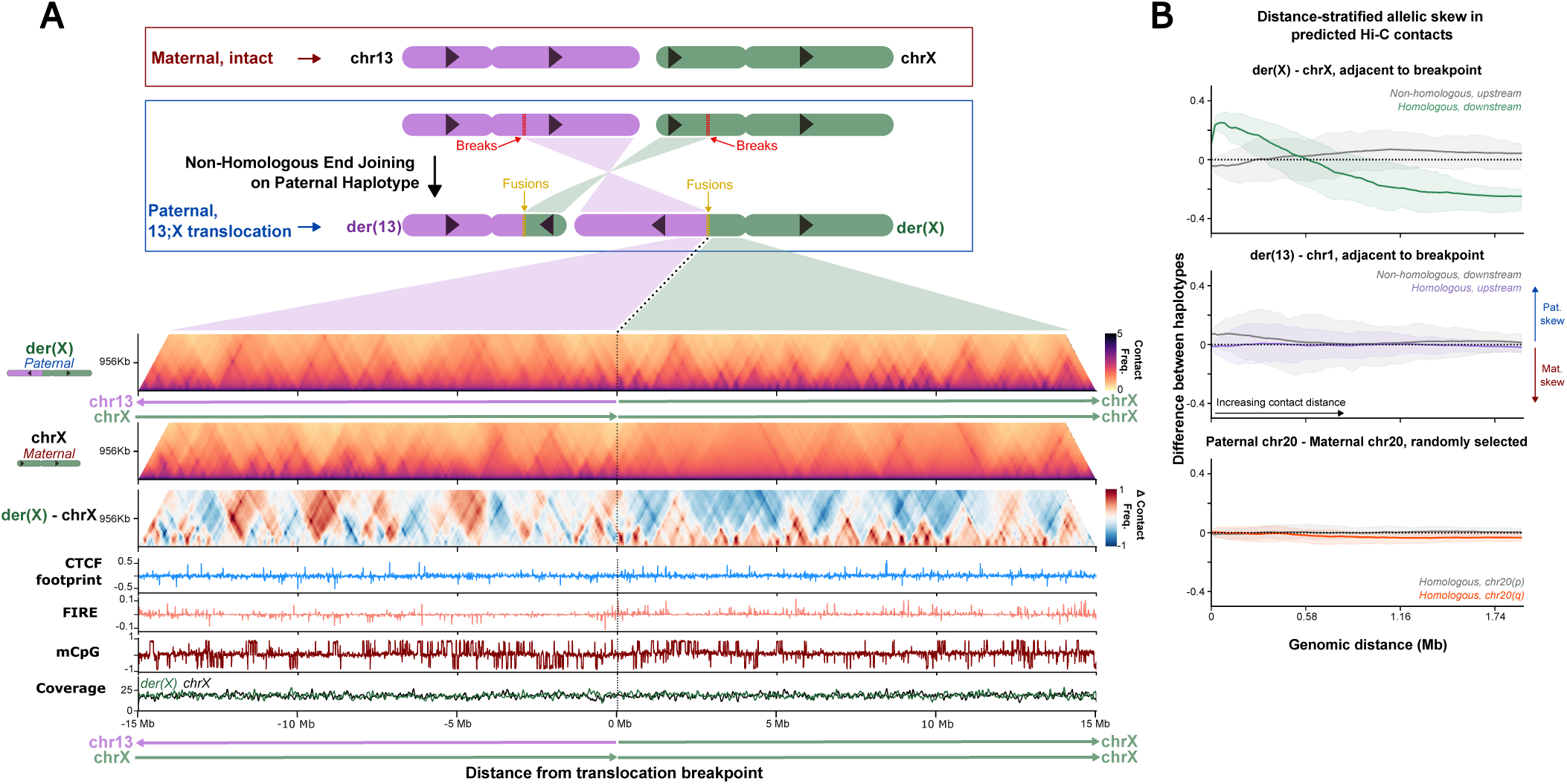
FiberFold predicts 3D genome organization in a clinically relevant 13;X balanced translocation. **A,** (Top) Cartoon depicting chr13-chrX balanced translocation affecting a 9-month-old female patient ^31^. (Middle) 30 Mb window depicting Hi-C predictions from FiberFold in the paternal derived chromosome (der(X)), maternal intact (chrX), and difference between them. Translocation position indicated by dotted line. Arrows denoting sequence origins and directionality. (Bottom) Input features for genomic region into FiberFold, and coverage. **B,** Mean distance-stratified difference per contig ± 8Mb, partitioned by breakpoint (except for 20q/p). Shaded region representing 25th and 75th percentile range. Labels indicate sequence of origin relative to the breakpoint and the homology between the sequences.

To test whether this is specific to X-derived chromatin, we also compared the homologous sequences of der(13) and chr13. There was a marked reduction in the magnitude of variability of haplotype-specific 3D chromatin structure between der(13) and chr13 (Fig. 5B, Supp. Fig. 5A,C,D). In comparison, the non-homologous sequences across the translocation breakpoint exhibited a similar pattern to that observed at the non-homologous regions between der(X) and chrX (Fig. 5B, Supp. Fig. 5C). In contrast, analyzing haplotype-specific 3D chromatin structure differences between the homologous p and q arms of chr20 revealed minimal pronounced variation (Fig. 5B, Supp. Fig. 5B,D).

Furthermore, FiberFold provided mechanistic insights into how der(13) alters the regulation of *PDK3*. Specifically, we previously observed a 3C interaction between a *MAB21L1* enhancer-like element (typically located on chrX) and the *PDK3* promoter (normally located on chr13) that was created selectively along der(13) ^31^. FiberFold predicts that this translocation results in the formation of a cross-breakpoint TAD along der(13) that spans 400–500 kb, encompassing and increasing the proximity between these genetic elements (Supp. Fig. 5E). In addition, we predict that this translocation also results in the formation of a cross-breakpoint TAD along der(X). (Supp. Fig. 5E). Together, these findings indicate that multiple genes along der(13) and der(X) are potentially undergoing loss-of-expression or ectopic gains-of-expression as a result this single balanced translocation.

## DISCUSSION

Leveraging advances in deep learning and single-molecule sequencing techniques, FiberFold enables researchers to use a single sequencing assay to assemble diploid genomes while building haplotype-specific maps of chromatin accessibility, CTCF binding, CpG methylation state, and 3D genome organization. FiberFold intentionally does not utilize genomic sequence in the underlying model, with the aim of increasing the generalizability of *de-novo* and haplotype specific Hi-C predictions. We observe accuracies similar to state-of-the-art, sequence-based models, indicating that chromatin accessibility, CTCF binding, and CpG methylation alone are sufficient to predict 3D chromatin structure at large scales. Furthermore, we demonstrate that this training strategy enables FiberFold to generalize beyond the cell type utilized for model training.

We predict widespread attenuation of TAD structures on the inactive X chromosome compared to the active X chromosome in humans. The distinctive 3D organization of the active and inactive X chromosomes has been recognized in mice crosses for over a decade ^64–69^, but this has been challenging to demonstrate in humans owing to the paucity of heterozygous variation along human chrX. Using two separate human cell lines with allelically skewed XCI, we demonstrate substantial differences between the 3-D organization of the Xi and Xa, providing a path forward for studying the 3-D chromatin organization of chrX in humans and other species with a lack of heterozygous variation along chrX. Notably, although there is substantial attenuation of TADs on the Xi, there are still many visually distinct domains forming (Figure 4D). This differs substantially from what has been previously reported in mice, where the inactive chrX adopts a bipartite structure consisting of 2 mega-domains and is nearly entirely devoid of TADs ^68–70^, indicating that the mechanistic details underlying XCI may diverge between mice and humans.

FiberFold has the potential to significantly streamline multi-omic analyses by enabling the joint quantification of genetic variation, CpG methylation, chromatin accessibility, protein occupancy, and 3D genomic structure in one assay (*i.e.,* Fiber-seq). The ability of FiberFold to generate haplotype-resolved Hi-C maps further adds to the potential of Fiber-seq as a powerful tool for uncovering the genetic and molecular basis of rare diseases. Specifically, we demonstrate that FiberFold can uncover patient-specific TAD alterations that are known to contribute to disease phenotypes. Consequently, we anticipate that applying FiberFold to patient-derived Fiber-seq data will enable the prioritization of genetic variants and elements that may be clinically impactful.

Recently, a scATAC-seq based model has demonstrated robust performance metrics by including co-accessibility matrices ^43^. Using single-molecule sequencing, we hope to incorporate similar signals to increase our model performance. FiberFold is currently limited in its ability to predict 3D genome structure across various window sizes and at various resolutions. We anticipate that the utilization of higher resolution micro-C data in our training, in combination with additional single-molecule Fiber-seq-derived features, will achieve even higher resolution and context lengths. Finally, as demonstrated using the Fiber-seq data from UDN318336, we anticipate that the utilization of donor-specific “isogenomic” ^71^ assemblies will enable the interrogation of the biological and clinical impacts of complex genomic variants that are not easily represented in reference assemblies.

## DATA & CODE ACCESSIBILITY

GM12878 and K562 Hi-C data was accessed with GEO accession: GSE63525. ENCODE ATAC-seq bigWig accession: ENCFF603BJO. CTCF ChIP-seq bigWig accession: ENCSR000DRZ. 30x GM12878 Fiber-seq data (training set) is available under NCBI Bioproject accession <u>PRJNA1124997</u>. This data can be interactively explored in the UCSC genome browser with hub: https://s3-us-west-1.amazonaws.com/stergachis-manuscript-data/2023/Vollger_et_al_long-read_multi-ome/GM12878_pacbiome/trackHub/hub.txt K562 Fiber-seq data is available under NCBI Bioproject accession PRJNA956114. This data can be interactively explored in the UCSC genome browser with hub: https://s3-us-west-1.amazonaws.com/stergachis-manuscript-data/2024/Vollger_et_al/FIRE/K562-PS00075/trackHub/hub.txt. Bottlenecked, phased GM12878 data can be accessed at https://s3-us-west-1.amazonaws.com/stergachis-manuscript-data/2024/Vollger_et_al/FIRE/GM12878/fire/GM12878.fire.bam. This data can be interactively explored in the UCSC genome browser with hub https://s3-us-west-1.amazonaws.com/stergachis-manuscript-data/2024/Vollger_et_al/FIRE/GM12878/trackHub/hub.txt. Training and prediction code is available at https://github.com/altemoselab/fiberFold.

## ACKNOWLEDGEMENTS

We would like to thank Professor Anshul Kundaje and all members of the Altemose Lab for helpful feedback.

## AUTHOR CONTRIBUTIONS

DD and NA conceived the project. AK and DD processed Hi-C data. DD designed, applied and analyzed the model and all results. JMF and CC assisted in data curation. MV, SN, and ABS provided Fiber-seq data and advised in Fiber-seq analysis of bottlenecked GM12878 and translocation patient data. MV and SN performed patient contig assembly. DD generated figures. NA and DD wrote the manuscript. All authors assisted in editing the manuscript.

## METHODS

### Hi-C Data

We used publicly available Hi-C data from two human cell lines: GM12878 and K562 (GEO accession number: GSE63525, SRA: SRP050102, enzyme: MboI), and processed them using HiC-Pro ^55,72,73^. The reads were aligned to the hg38 reference genome, and each sample was processed independently. Following alignment, validated interaction pairs for each cell line were merged to generate a combined contact matrix. We applied Iterative Correction (ICE) normalization to the matrices using HiC-Pro’s built-in functions. After normalization, the .matrix files were converted to the cooler format^74^ using the cooltools package^75^. Subsequently, the .cool files were transformed into .npz format to be used as input for further analysis (cool2npy.py script from C.Origami^38^).

### ATAC-seq and ChIP-seq Data

We used publicly available ATAC-seq and ChIP-seq data from the ENCODE consortium^57–59^. ATAC-seq for figure 1A tracks was downloaded from experiment accession ENCSR637XSC with bigWig file accession ENCFF603BJO. For CTCF ChIP-seq data we accessed the bigWig file accession ENCFF644EEX under experiment accession ENCSR000DRZ. For Rad21, ChIP-seq data we used the bigWig file with accession ENCFF571ZJJ, under experiment accession ENCSR000EAC.

### Fiber-seq Data and Processing

We used publically available 30x Coverage GM12878 Fiber-seq data^31^. Deep, haplotype phased GM12878 data was obtained from ref. 27. K562 Data was obtained from ref. 72. We used publically available FIRE tracks for all GM12878 data. K562 FiRE peaks were called using fibertools fire and fibertools pileup with default parameters^76^. To identify CpG we first used pbjasmine to call 5mC instances on single-molecules using parameters --keep-kinetics --min-passes 3. We then used pb-CpG-tools (https://github.com/PacificBiosciences/pb-CpG-tools) to generate one-dimensional bedGraph style tracks for training with parameters --modsites-mode reference and –model pileup_calling_model.v1.tflite (model available on the above git page).

### Calling CTCF Peaks and CTCF Direction

We downloaded CTCF sites (Jaspar Motif ID MA0139.1) in the hg38 genome from Jaspar (http://expdata.cmmt.ubc.ca/JASPAR/downloads/UCSC_tracks/2022/hg38/MA0139.1.tsv.gz) ^77^. For sites where there was more than one instance of a predicted CTCF motif, we selected the motif with the higher score to be the representative motif.

We defined 3 individual modules from positions 0-6,6-13, and 13-19 (0 based, half open) (https://fiberseq.github.io/fibertools/extracting/footprint.html) within the representative motif and used *fibertools footprint* with default parameters to perform footprinting.

To convert these single-molecule footprint calls into a continuous bigWig style track for prediction and training we then use the encodeFP.py script to process the bit flag encoding the single-molecule footprint code. Briefly, using the output of ft footprint, we iterate through every motif in the genome and at every instance of a fiber-seq read overlapping that motif, we check that all these conditions are satisfied:

1. The motif does not lie within a nucleosome
2. All modules containing an Adenine or Thymine are footprinted

If these conditions are satisfied then we classify that motif-molecule overlap instance as a footprint. We then divide the number of footprints by sequencing depth at the motif to quantify the CTCF footprinting score. To encode CTCF motif direction we generate a bigWig file where CTCF motif instances in the sense orientation are represented with 1 and instances in the antisense orientation are represented with -1.

### Filtering Low-Quality Regions

Regions of the genome that are poorly mappable, low-coverage, or noisy were identified and filtered from analysis. The problematic regions included telomeric regions (50 kb from each chromosome end), centromeric regions (from the UCSC Genome Browser, hg38 assembly, centromeres track), the ENCODE blacklist ^78^, and regions with low coverage in GM12878 fiber-seq data.

To aggregate problematic areas, we merged regions that were within 2.5 kb of each other. Additionally, regions shorter than 5 kb in length were excluded from the blacklist, yielding a final set of regions to be ignored during training. Any training window that overlapped these problematic areas by more than 50 kb was filtered out, and the resulting filtered dataset was then segmented into train, test and validation sets.

### Model Architecture

Model architecture and the training regiment was adapted from C.Origami^38^ with a few notable modifications. First, we do not include DNA sequence information. Second, we include CpG methylation as a track for training. Third, we include CTCF direction as input to the model. This results in a 50% reduction in parameter size in the initial encoder block of the model, as we do not include an explicit sequence encoder, just an epigenomic encoder. All Hi-C data is log transformed. All FiRE/CpG/CTCF track data is depth-normalized and thus represented between 0-1. The transformer and decoder architecture from the C.Origami model are unchanged. Similar to other 3D-genome prediction models, we optimized against a MSE loss function comparing the predicted Hi-C map with the experimental Hi-C map.

### Model Training

Chromosomes 10 and 15 were held out for validation and testing, respectively. For the remaining chromosomes we apply a sliding window with a step of 36 kb to generate a list of windows that can be quickly sampled during training. We then filter out low-quality windows from training as defined above with bedtools. During training we apply a series of data augmentations to each window described below:

1. In training we apply a random shift to every training window ( maximum shift +/ 10 kb ) every epoch.
2. Like C.Origami^38^, we apply a gaussian shift to all features in every window with mean 0 and standard deviation of 0.1.
3. We reverse the Hi-C matrix and the feature matrix with 50% probability.

We trained the model on 4 A100 GPUs (40 Gb) on Stanford’s Sherlock cluster with an Adam optimizer, a learning rate of .0004, and a batch size of 8. We trained until model performance did not improve for 15 epoch iterations.

### Correlation Calculations

To calculate bulk map-wide experiment-prediction correlations across the test chromosome (chr15) we use scipy’s pearson or spearman function to calculate the correlations between the experimental or predicted map. For pixel-to-pixel comparisons we compare all pixels against each other that are at least 16,384 bp away from each other, or simply an offset of 2 onward from the diagonal. To calculate insulation across windows we applied the same algorithm used in Tan et. al. 2024^38^. Insulation was calculated with a radius of 500 kb.

### Cross Cell-Type Performance

To calculate model performance in K562 cells, we predicted genome-wide Hi-C maps using publicly available K562 Fiber-seq data. We employed a sliding window approach with a step size of 216 kb, resulting in the prediction of 2968 Hi-C maps for K562 cells. For evaluation, we quantified both map-wide correlations and pixel-by-pixel correlations, similar to our previous approach with GM12878 cells.

For comparisons between GM12878 and K562 Hi-C maps, we normalized for differences in sequencing depth between the samples. This normalization was done by calculating the mean sequencing depth across the corresponding diagonals of the Hi-C maps, determining the ratio between the cell lines, and adjusting the K562 contact information by multiplying its array by this ratio.

### Phasing Fiber-seq Information

Phased, bottlenecked GM12878 Fiber-seq data was separated into bam files associated with the respective haplotypes using samtools. FiRE, mCpG, and CTCF footprinting tracks were then generated identically as above for each haplotype. The data from each haplotype were then processed separately.

### Haplotype-Specific Predictions in GM12878

To analyze haplotype-specific contact maps genome-wide, we generated 2 MB windows across the genome with a step size of 500 kb. Using a previously described blacklist, we filtered out all low-quality windows. Additionally, to avoid artifacts from reference mapping due to large structural variants, we excluded windows where either haplotype had more than 10 sites with pixel values equal to or less than 0, indicative of localized mapping errors. After applying these filters, we retained 3441 sites for comparison.

### Quantifying Haplotype-Specific Contribution to Experimental Signal

To calculate the relative contribution of haplotype-specific contacts to observed experimental signals, we first separated chromosome X-specific pixels from autosome-specific pixels. For each group, we then binned the experimentally derived pixel values into decile groups. For each decile bin, we selected all predicted pixels from both haplotypes that corresponded to the pixels within that bin and calculated the 5th and 95th percentiles for these pixels. Finally, we determined the proportion of predicted pixel values from each haplotype that fell below the 10th percentile or above the 90th percentile, respectively.

### Generating Maps Longer Than 2 MB

To generate maps larger than 2 MB, we stitched together contiguous overlapping maps. We used a step size of either 131,072 bp (for autosome-X translocation) or 262,144 bp (for GM12878). Each individual Hi-C map was then rotated 45° counterclockwise to align the inter-bin contact diagonal with the x-axis. Subsequently, we appended each overlapping map to a 3-dimensional array. At any overlapping site, we calculated the mean contact frequency across all maps that contained predictions at that site. To ensure contiguous information, we trimmed the maps, removing either 16 or 32 pixels from the top, corresponding to the 131,072 bp or 262,144 bp step, respectively. This process resulted in long-distance stitched Hi-C maps that can represent 3D genomic contacts at a maximum distance of 1.967 MB or 1.835 MB.

### Translocation Patient Genome Assembly and Contig Classification

To generate a contig-level assembly for the patient with the autosome-X translocation, we first combined all long-read Fiber-seq data from the proband into a single FASTA file. We then constructed a k-mer hash table using parental short-read data via yak v0.1 with the following command: *yak count - b 37*. Contigs were generated using hifiasm^79^ v0.20.0 in trio mode with the command: *hifiasm fiber_seq.reads.fa -1 paternal.yak -2 maternal.yak*. Next, we aligned the assembled and phased contigs to the hg38 reference genome using minimap2 in assembly mode with the command: *minimap2 -ax asm5 --MD --eqx -c* ^80^. We identified the location of the previously known breakpoint ^31^ in the contigs using the liftover utility from rustybam. We then manually inspected all contigs overlapping the identified breakpoint within IGV, focusing on contigs longer than 10 MB and assessing their continuity on both sides of the breakpoint. Maternally derived contigs were expected to show continuity for both chrX and chr13 sequences on either side of the breakpoint. Conversely, derivative chromosome contigs, der(X) and der(13), should demonstrate continuity for the original chrX or chr13 sequence on one side of the breakpoint and exhibit continuity for the translocated sequence in an inverted orientation on the opposite side, resulting in an alternate strand alignment beyond the breakpoint.

### Haplotype-Specific Hi-C Predictions in the Autosome-X Translocation Patient

We then mapped the proband Fiber-seq data back onto the assembled and phased contigs and calculated the FiREs, CTCF footprinting score, and CTCF motif orientation, following the same procedures as before. Due to the unpolished nature of the assembly, we used the parameter *-q 0* in the pb-CpG-tools command when computing CpG methylation tracks to include unphased methylation data as well.

### Average Distance-Stratified Differences Between Haplotypes

To calculate the distance-stratified differences between haplotypes in the autosome-X translocation patient, we first generated a contiguous 16 Mb Hi-C map for each contig, centering the map at the breakpoint location on that contig. We then calculated the differences between these maps for the two haplotypes, using the breakpoint position as the boundary to partition the comparison into two groups. Subsequently, we calculated the mean difference at every length scale, as well as the 25th and 75th percentiles, and the standard deviation of the difference. Using the partitioned data, we then calculated distance-stratified p-values for differences between the two groups across all data points at every distance with a two-sided Mann-Whitney U test.

## SUPPLEMENTARY FIGURES

**Supplementary Figure S1:**
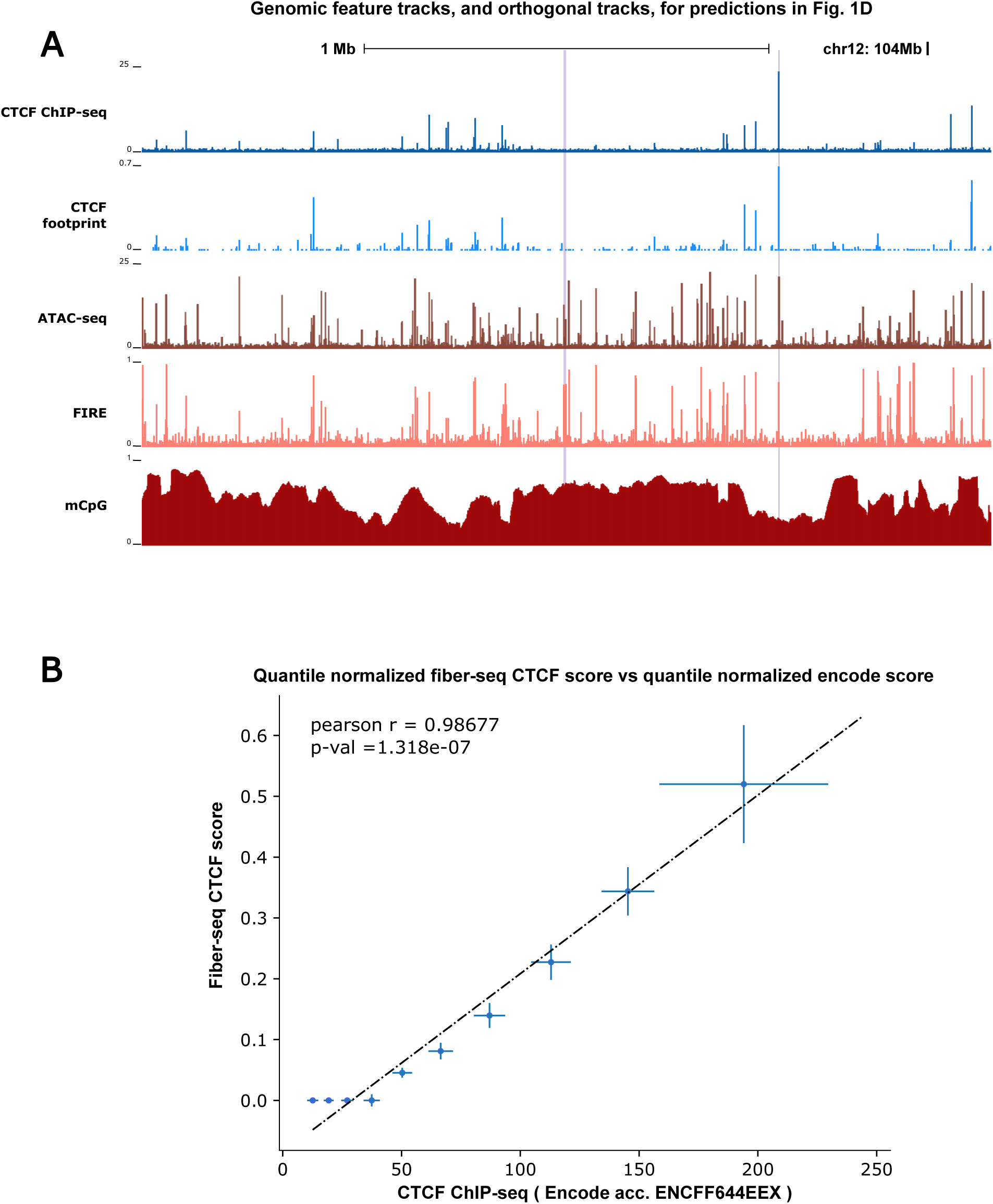
Fiber-seq capture both large and fine scaled features impacing 3D chromatin structure. **A,** Comparison of binned CTCF ChIP-seq data against binned Fiber-seq CTCF footprinting scores. **B**, Decile binned CTCF ChIP-seq signal and CTCF footprinting score as determined by Fiber-seq. Points represent median values, bars represent standard deviation of values. Points denote median values, error bars represent standard deviation. Dotted line represents regression line.

**Supplementary Figure S2:**
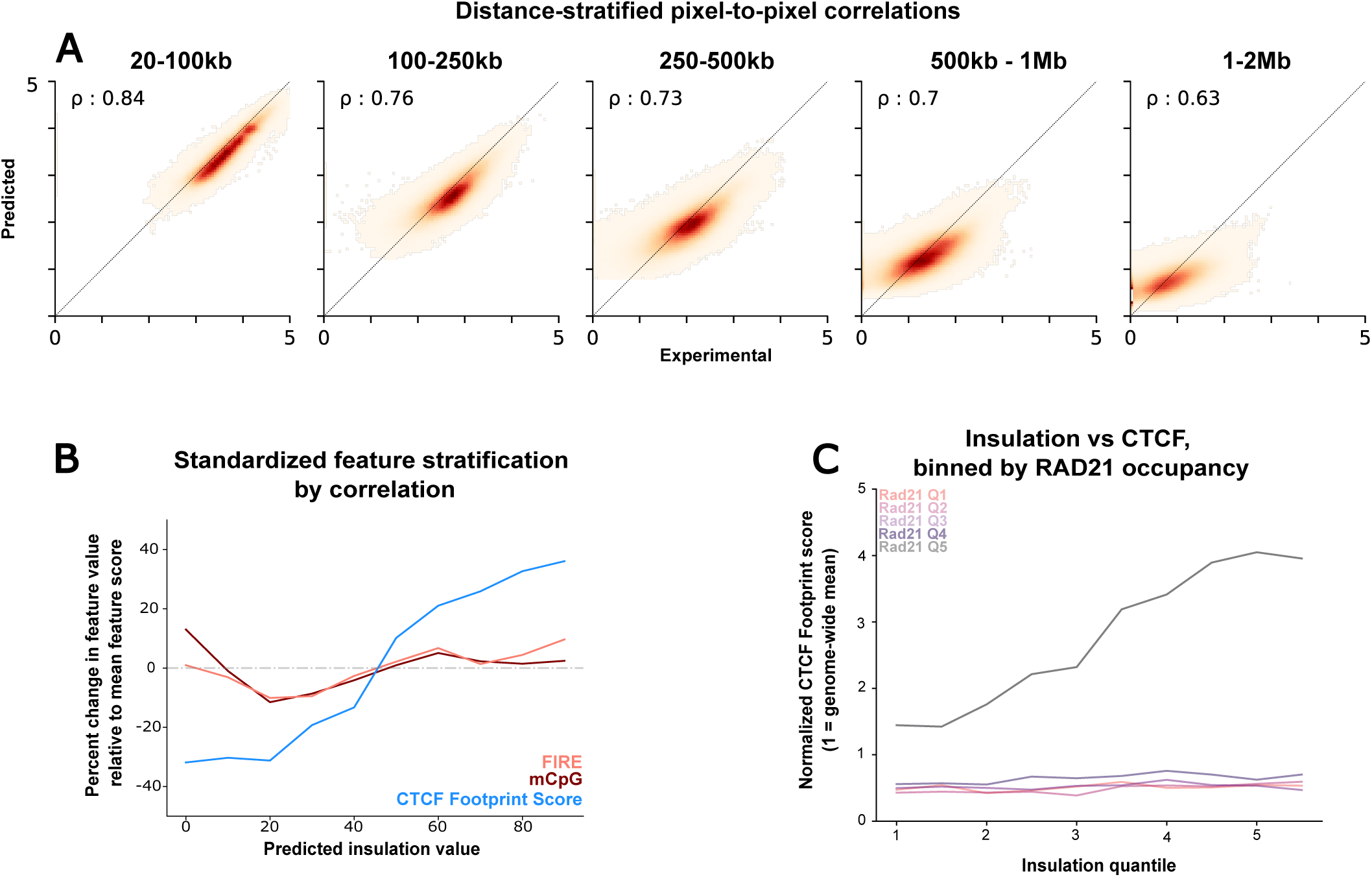
Distance and Feature Stratification of FiberFold performance in GM12878. **A,** Experimental and predicted pixel correlation as displayed in Fig.1C stratified by genomic distance for chr15 (held out test set) in GM12878. Pearson R displayed in upper left corner. **B,** Feature stratification in test-set chr15, binned by predicted insulation value. Features are standardized by dividing by the global feature mean (chr15). **C,** CTCF footprint score derived by Fiber-seq, binned by insulation and stratified by overlapping Rad21 ChIP-seq signal.

**Supplementary Figure S3:**
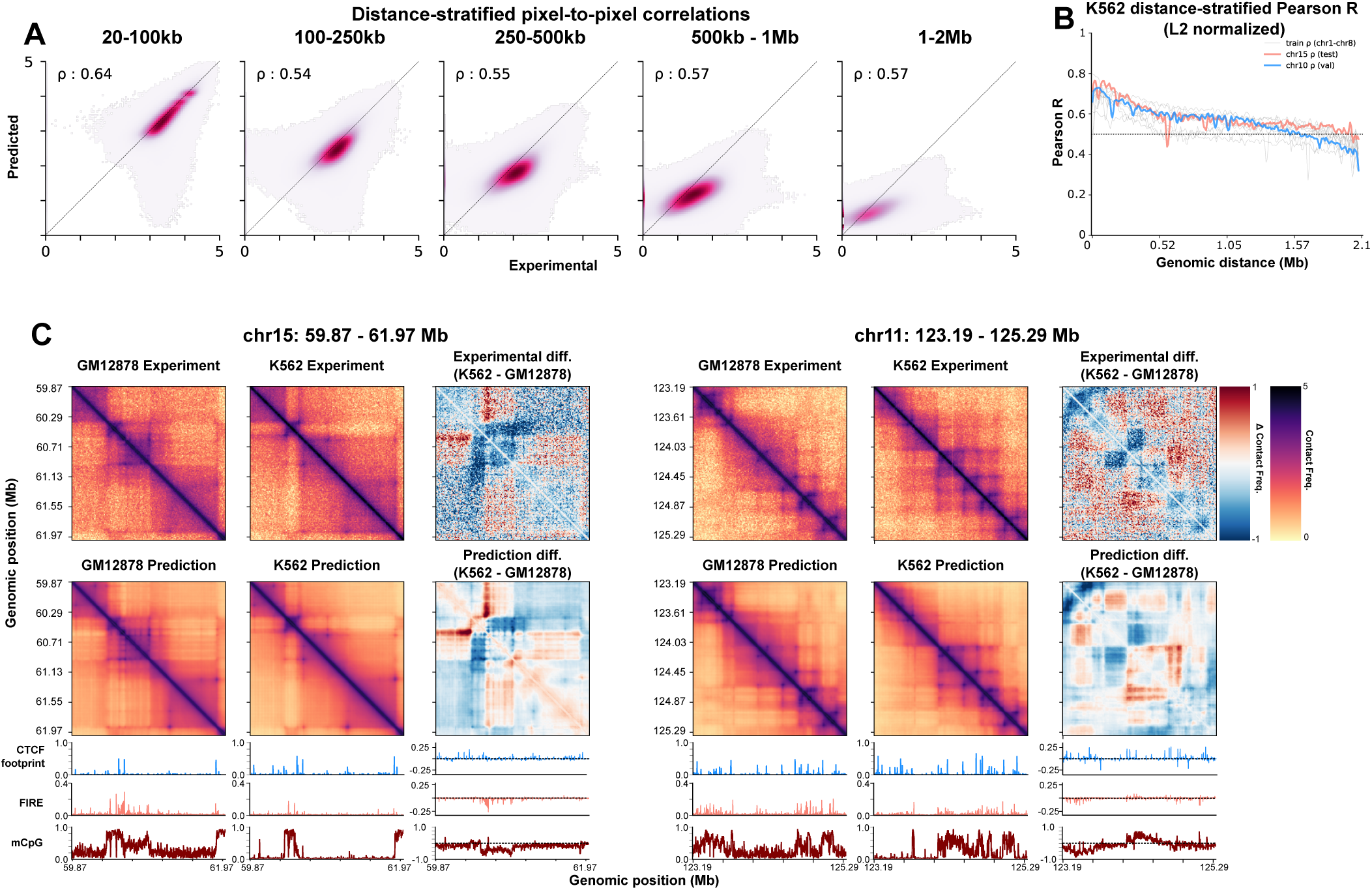
FiberFold performance in a *de novo* cell line (K562) **A,** Experimental vs. Pixel value across all chromosomes for K562, stratified by distance. **B,** Pearson R across all chromosomes in K562, stratified by distance. **C,** Examples of Experimental and Predicted Hi-C data for K562 (middle), GM12878 (left), and the difference between them (right). Underlaid are the input features (left, middle) and the input feature difference (right)

**Supplementary Figure S4:**
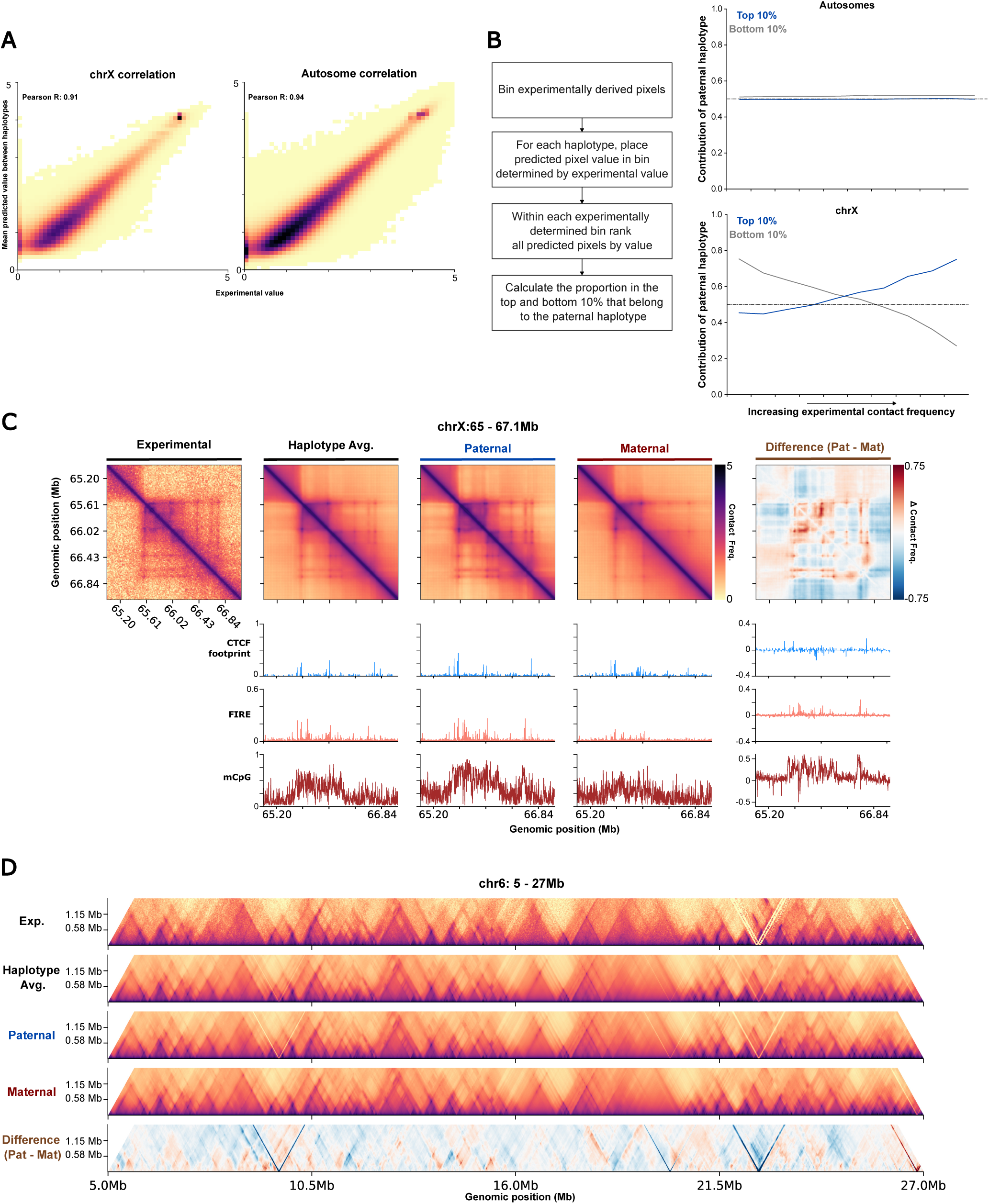
Attenuated Maternal TADs on chrX. **A,** Experimental Pixel vs. Haplotype-average predicted Pixel Value in chrX and autosomes. **B,** Paternal haplotype contribution to top and bottom 10% of aggregated prediction signal, stratified by experimental pixel value in chrX and autosomes (Methods). **C** chrX example of haplotype-specific prediction differences in Hi-C maps. **D,** Haplotype-specific differences displayed across a randomly selected 22 Mb window in chr6.

**Supplementary Figure S5:**
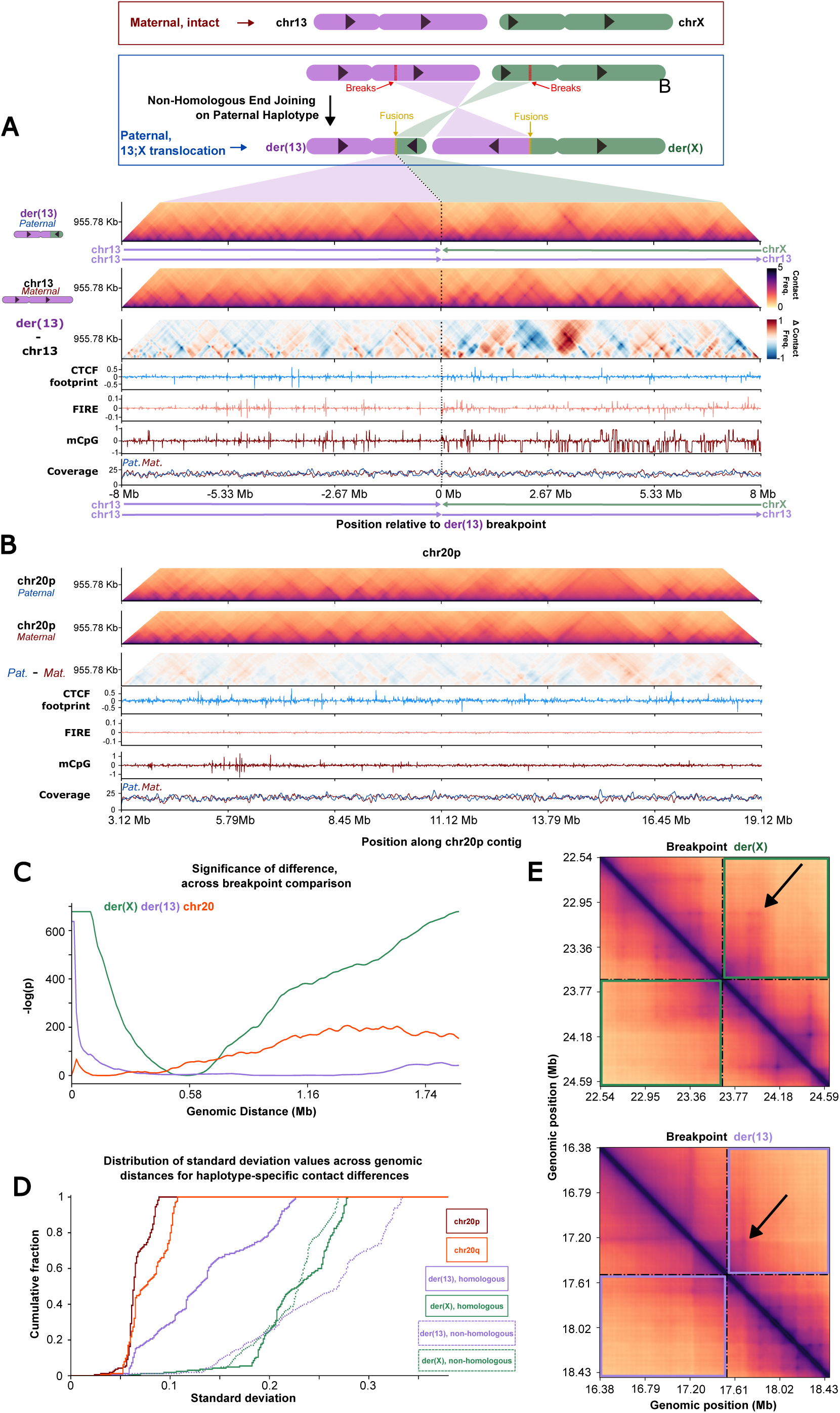
Distribution of 3D chromatin structures around der(13) and der(X) **A,** (Top) Cartoon depicting chr13-chrX balanced translocation affecting a 9-month-old female patient ^31^. (Middle) 16 Mb window depicting Hi-C predictions from FiberFold in the derived chromosome (der(13)), intact (chr13), and difference between them. Translocation position indicated by dotted line. Arrows denoting sequence origins and directionality. (Bottom) Input features for genomic region into FiberFold, and coverage. **B,** Hi-C maps displayed as in A, for contiguous regions on haplotype-specific contigs representing chr20p in the patient, and the difference in their 3D contacts. **C,** Distance stratified significance values (Mann Whitney) for predicted contact differences in der(X), der(13), and chr20. **D**, Cumulative distribution of the standard deviation in contact differences across genomic distances for each comparison group. Solid lines indicate comparisons between homologous sequences (chr20p, chr20q, region of homology upstream of breakpoint on der(13), region of homology downstream of breakpoint on der(X), non-homologous region downstream of breakpoint on der(13), and non-homologous region upstream of breakpoint on der(X). **E,** FiberFold-predicted Hi-C maps at breakpoints on derived chromosomes (dashed line). Colored boxes indicate region representing cross breakpoint contacts. Arrows denote regions with increased 3D contact frequency across breakpoint.

